# Interactions between *Streptococcus agalactiae* and *Candida albicans* affect persistence and virulence

**DOI:** 10.1101/2025.10.07.681024

**Authors:** Kathryn Patenaude, Chloe N. Bossow, Anna Lane, Marc St-Pierre, Robert T. Wheeler, Melody N. Neely

## Abstract

*Streptococcus agalactiae* (Group B Streptococcus or GBS), a Gram-positive bacterium, and *Candida albicans*, a polymorphic fungus, are commensal microbes in most of the population they colonize. However, for certain patients they can cause severe and sometimes fatal infections. Previous research has indicated that GBS and *C. albicans* can synergize to enhance the colonization of GBS in the bladders of mice, but not much was known prior to this study about how interactions between GBS and *C. albicans* alter treatment effectiveness and infection outcome *in vivo*. Results showed that interactions between the two opportunistic pathogens were influenced by media nutrient availability, and that the presence of *C. albicans* in a culture reduces the effectiveness of certain antibiotics against GBS *in vitro.* This study also utilized a larval zebrafish model to investigate differences in virulence in solo infections vs co-infections with both pathogens *in vivo*. Co-infections of GBS and *C. albicans* into the otic vesicle were found to have increased virulence compared to solo infections of either pathogen. Co-infection also led to an increased GBS burden compared to solo GBS infections. Co-infections of GBS and *C. albicans* by yolk sac injection were not more virulent than solo infections with either pathogen. However, the antibiotic clindamycin was less effective in preventing mortality in co-infections compared to solo GBS infections. Overall, these findings highlight how interactions between GBS and *C. albicans* can influence treatment effectiveness and virulence during infection.

## Introduction

*Streptococcus agalactiae*, also known as Group B streptococcus or GBS, is a Gram-positive, opportunistic pathogen that colonizes the gastrointestinal and/or vaginal tract of roughly 20-30% of healthy people worldwide (1). While this microbe is a commensal in most of the population, certain patients are at a higher risk of serious infection by GBS, including the elderly, immunocompromised, patients suffering from chronic conditions like diabetes and cancer, pregnant women, and newborns (2–5). GBS colonization of the vaginal tract in pregnant women is common, with 10-30% of pregnant women estimated to be colonized (6). This colonization is a major risk factor for invasive GBS disease in neonates if exposed to the pathogen in utero through ascending infection or at the time of delivery. Newborns exposed to GBS at time of birth are at risk of contracting serious, and sometimes fatal, early-onset (EOD) or late-onset (LOD) bacterial meningitis infections, with surviving children facing life-long neurological complications (6–9). There is no commercially available vaccine for GBS, and there are very limited treatment options for mothers colonized with GBS prior to or during delivery (10, 11). To treat pregnant mothers colonized with GBS, intrapartum antibiotic prophylaxis (IAP) is administered directly before and during delivery (6). While IAP has been shown to be effective at reducing the rates of EOD clinically, the consequences from this treatment, including the development of antibiotic resistant GBS strains and disruptions to the microbial flora of both the mother and their offspring, are not well understood (6, 12–15).

Historically, research on an infectious pathogen is conducted by investigating the pathogen in solo. However, many tissue environments are microbially diverse, unique to each patient, and can be rapidly altered depending on environmental conditions. Polymicrobial interactions have been shown to directly influence treatment effectiveness *in vitro*, as well as affect treatment and infection outcome in patients clinically (16–21). Polymicrobial interactions between GBS and other microbes co-colonizing the vaginal tract are poorly understood. While the most common microbes found to be colonizing the human vaginal tract belong to the *Lactobacillus* genus, other microbes can colonize the human vaginal tract and overgrow (sometimes leading to infection) following disruptions to the lactobacilli population, including *Candida albicans* (22, 23).

*C. albicans* is an opportunistic pathogenic yeast that colonizes the vaginal tract of ∼30% of women (24). Like GBS, *C. albicans* is a commensal in most of the population, but can cause severe infections in immunocompromised patients, which include pregnant women and newborns (25–28). Oropharyngeal candidiasis (thrush) caused by *C. albicans* can occur in newborns due to passage from a colonized mother to the newborn during delivery (29). *C. albicans* is also the leading cause for vulvovaginal candidiasis (VVC), with 75% of women having a VVC infection at least once in their lifetime, and ∼8% of those cases being recurrent (30, 31). Pregnant women are considered to be at a higher risk of developing VVC caused by *C. albicans* (32). Because *C. albicans* and GBS can colonize and infect the same types of patients and tissue environments, interactions between these microbes *in vivo* could influence factors like treatment effectiveness and infection outcome following challenges by these opportunistic pathogens.

The relationship between *C. albicans* and other streptococcal species has been well documented, as *C. albicans* has been shown to exhibit synergy with multiple oral streptococcal species, including *Streptococcus mutans* (33, 34), *Streptococcus oralis* (35, 36) and *Streptococcus gordonii* (37, 38) to enhance biofilm formation and virulence. While there has been less reported on the interactions between GBS and *C. albicans*, there is clinical evidence that *C. albicans* and GBS are often co-isolated together from colonized patients (39, 40). Previous research has indicated that co-association between GBS and *C. albicans* is modulated through hyphal-specific surface adhesion protein *Als3* (41), while another study indicated that GBS can prevent *C. albicans* hyphal development (42), indicating that interactions between these two pathogens are complex and likely dependent on environmental conditions. *C. albicans* can also promote bladder colonization by GBS, further establishing the role that interactions between the two organisms have in the carriage of both microbes (43). While previous studies show that these microbes can co-associate to enhance the colonization of both the bladder and vaginal tract (39, 40, 43), the role this co-association has on treatment effectiveness and virulence during infection has not been explored prior to this study.

This study demonstrated how co-association between GBS and *C. albicans* alters growth rate, treatment effectiveness and influences virulence during infection. Results demonstrated synergistic interactions during co-culturing of *C. albicans* and GBS. Interactions between *C. albicans* and GBS led to decreased susceptibility of GBS to the antibiotics erythromycin and clindamycin *in vitro* and reduced the effectiveness of clindamycin treatment against GBS infection *in vivo*. Results demonstrated a clear synergistic effect in survival outcomes of co-infections of GBS and *C. albicans* in comparison to solo infections by either pathogen utilizing a zebrafish infectious disease model. This research highlights how polymicrobial interactions can influence the treatment effectiveness and virulence of the pathogens GBS and *C. albicans* when co-colonized in comparison to solo colonization with either pathogen.

## Results

### GBS and *C. albicans* synergize to enhance growth *in vitro*

GBS and *C. albicans* are opportunistic pathogens that can adapt to colonize multiple sites in the body, including the vaginal tract. The vaginal tract can be a difficult environment for microbes to colonize due to low pH and limited nutritional availability, forcing microbes to adapt in order to survive (44, 45). To investigate the role nutrient availability plays in the co-association between GBS and *C. albicans,* these organisms were co-cultured together or grown separately in nutrient rich (Todd Hewitt Broth with 0.2% supplemented yeast, THY B) or nutrient poor (serum free RMPI 1640) media environments and monitored for growth over time. In RPMI media, the growth of GBS was significantly increased in co-cultures with *C. albicans* at both 6 hours (4.8 fold, *p=0.04), and at 24 hours (11.6 fold, *p=0.01) following initial culture compared to solo cultures of GBS (Figure 1A). *C. albicans* viability was not significantly different in solo or co-cultures with GBS at either 6 or 24 hours post culture in nutrient poor media (Figure 1B).

**Figure 1:**
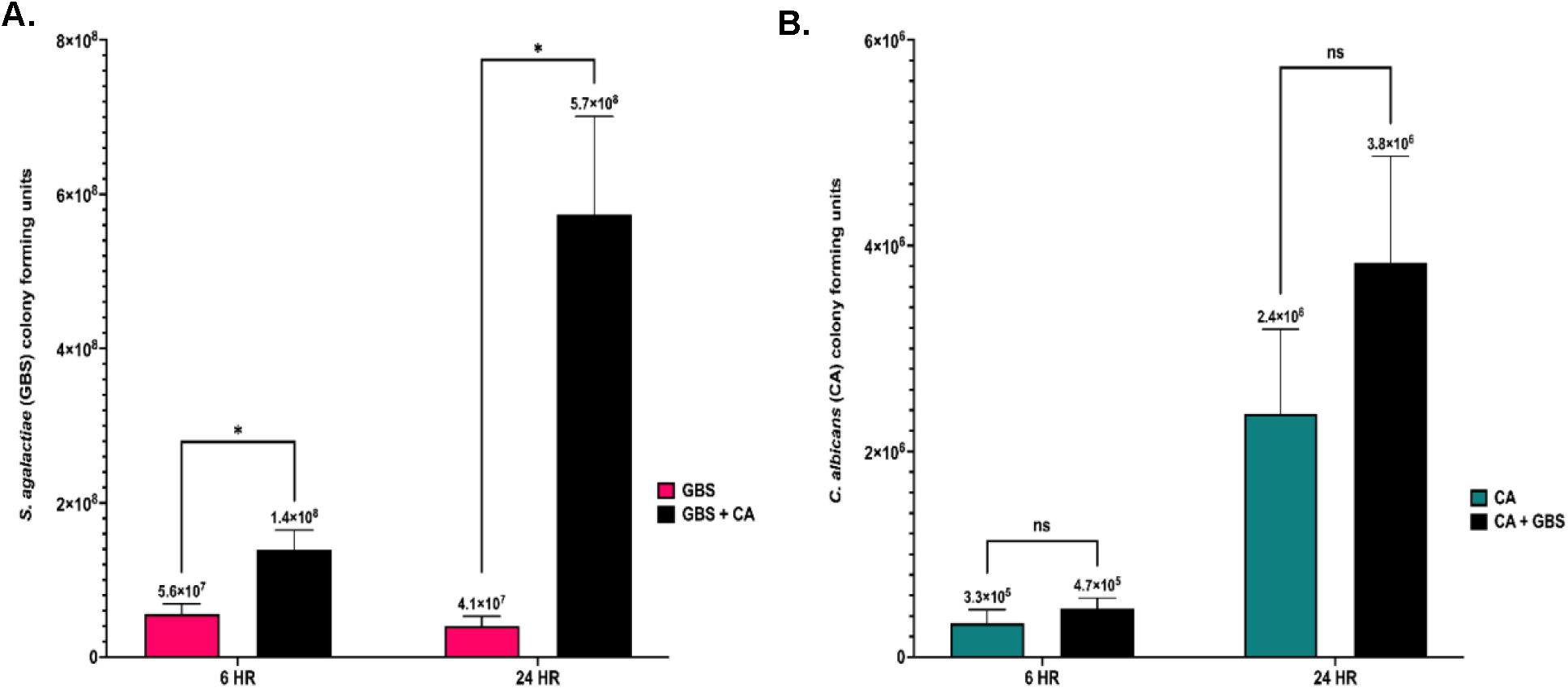
Interactions between GBS and *C. albican*s during co-culture in low nutrient media alters growth and viability. **A.** Colony growth of GBS following solo or co-culture with *C. albicans* at 6-and 24-hours post culture in nutrient poor media **B.** Colony growth of *C. albicans* following solo or co-culture with GBS at 6-and 24-hours post culture in nutrient poor media. Results are from combined experiments, and experiments were performed in triplicate. Statistical significance was calculated by unpaired two tailed student t-test, *p<0.05, **p<0.005, ****p<0.00005.

Alternatively, neither GBS or *C. albicans* cell growth was significantly different when in solo or co-cultures when grown in THY B (nutrient rich media) at 24 hours post culture (Supplemental figure 1). The increased growth of GBS in the presence of *C. albicans* in RPMI media is not strain specific, as this effect was also observed in co-cultures of *C. albicans* and COH1 WT (a serotype III GBS strain), indicating that the increase of GBS growth in the presence of *C. albicans* in nutrient poor environments may be conserved amongst GBS strains (Supplemental figure 2). However, using three additional *C. albicans* clinical vaginal isolates (46, 47) resulted in varying amounts of enhanced GBS growth (Supplemental figure 3). The ability of *C. albicans* to significantly increase the growth of GBS in RPMI media is dependent on the viability of *C. albicans* cells, as GBS growth in RPMI media with heat-killed *C. albicans* cells was not significantly different compared to solo GBS cultures following 24 hours of incubation (Figure 2). To investigate early interactions between GBS and *C. albicans* in co-culture environments in RPMI media, GBS and *C. albicans* were cultured both in solo and in co-cultures, and cell growth was measured every 2 hours for a 6-hour time period. GBS growth was significantly increased by the presence of *C. albicans* compared to solo cultures as early as 4 hours post co-culture (7.2-fold, **p=0.0010) (Figure 3A). Notably, fewer *C. albicans* cells were recovered in RPMI media at 2 hours and 4 hours post-inoculation compared to the original starting inoculum (Figure 3B). The presence of GBS in co-cultures with *C. albicans* reduced the loss of *C. albicans* cells in comparison to solo cultures of *C. albicans* at both 2 hours (5.7-fold, **p=0.0051) and 4 hours (5.1-fold, *p=0.0144) co-culture, indicating that the presence of GBS may be beneficial for the recovery of *C. albicans* cells in RPMI media (Figure 3B).

**Figure 2:**
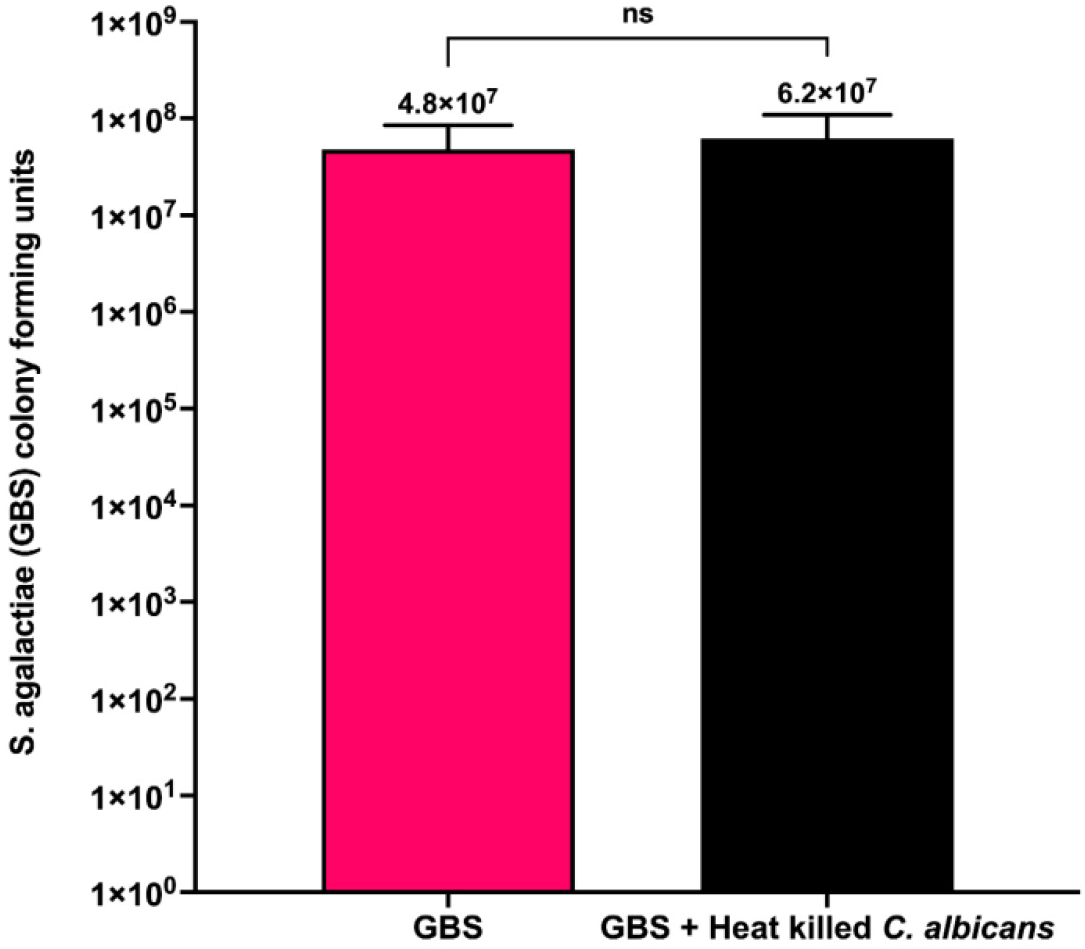
*C. albicans* must be viable to cause an increase in GBS growth during co-culture. Growth of GBS alone or in co-culture with heat-killed *C. albicans* cells in nutrient poor media for 24 hours. Results above are from combined experiments, and experiments were replicated 4 times (n=4). Statistical significance was calculated using an unpaired two-tailed student’s t test.

**Figure 3:**
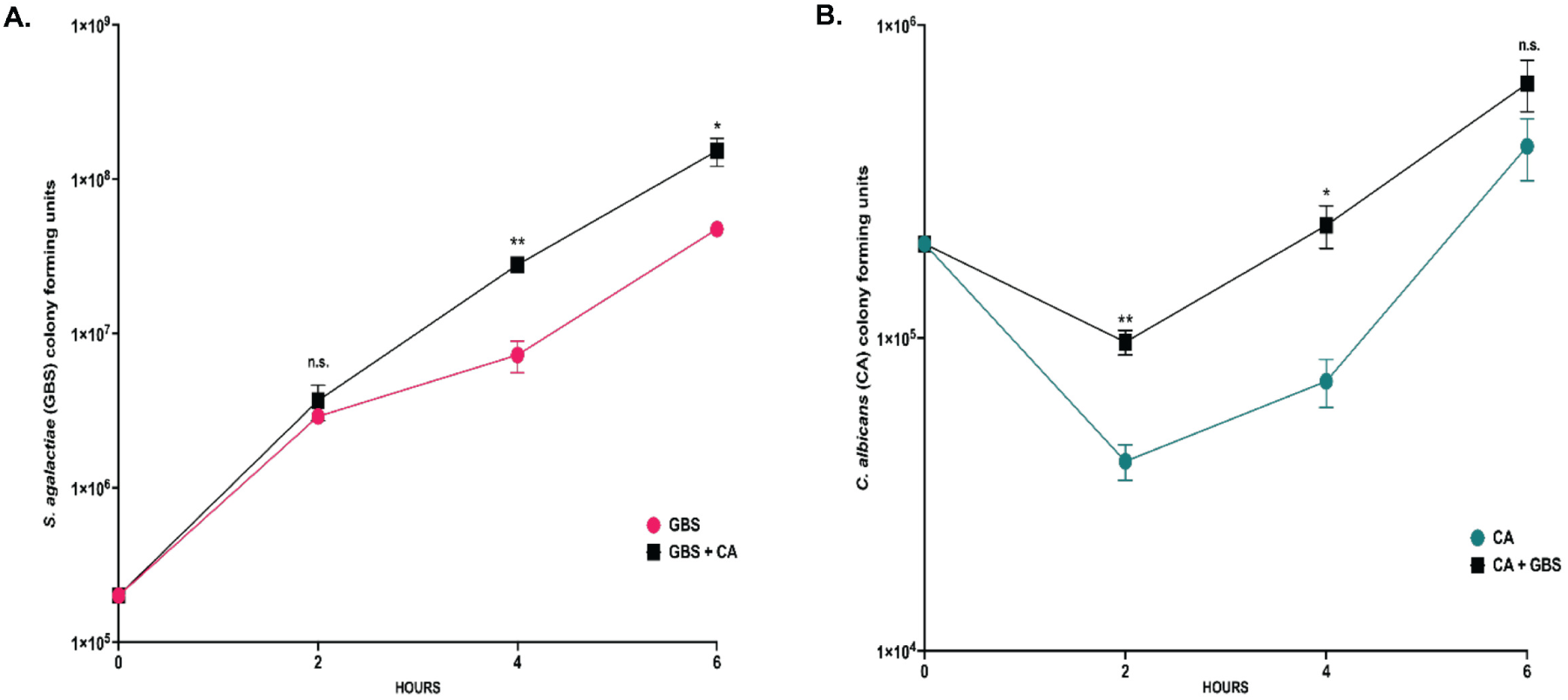
The growth rate of GBS and *C. albicans* in solo and co-cultures. **A.** Concentration of GBS cells (colony forming units) in solo (magenta) and co-cultures with *C. albicans* (black) in nutrient poor media. **B.** Concentration of *C. albicans* cells (colony forming units) in solo (blue) and co-cultures with GBS (black) in nutrient poor media. Results are from combined experiments, and experiments were performed in triplicate and statistical significance was calculated by unpaired two tailed student t-test for each time point, *p<0.05, **p<0.005, ****p<0.00005.

### *C. albicans* hyphal formation is not required for increased growth of GBS

*C. albicans* is a polymorphic microbe that can proliferate as yeast, pseudohyphal, or hyphal cells depending on environmental conditions. Research investigating the role of *C. albicans* hyphae during interactions with GBS have shown conflicting results, as it has been shown that certain GBS strains may inhibit hyphal growth by *C. albicans*, while others have shown that GBS can actually benefit from attaching to hyphae *in vitro* (41, 42). To investigate the role of hyphae in the increased growth of GBS observed in the presence of *C. albicans* in RPMI media, GBS was co-cultured with either a strain of *C. albicans* that has overexpression of a repressor of hyphal formation and therefore remains in its yeast morphology *in vitro* (*NRG1^OEX^-iRFP*) or the parent strain (CAF2-FR) (48–51). Fluorescent images of co-cultures of GBS and the parent strain CAF2-FR, compared to growth of CAF2-FR alone, following a 24-hour incubation in RPMI media, demonstrated that CAF2-FR cells produced hyphae in the nutrient poor media (Figure 4A), while *NRG1^OEX^-iRFP* cells were unable to do so and remained in a yeast morphology (Figure 4B). The presence of GBS did not alter the morphology of *NRG1^OEX^-iRFP* or CAF2-FR (Figure 4C & 4D), indicating that the presence of GBS did not prevent CAF2-FR from forming hyphae in RPMI media (Figure 4C, Supplemental Figure 4). The ability to form hyphae was not required for *C. albicans* to significantly increase the growth rate of GBS in RPMI media, as *C. albicans* strains CAF2-FR and yeast locked *NRG1^OEX^-iRFP* caused significantly (p=0.001 & p=0.0008, respectively) increased growth of GBS (Figure 4E) following a 24-hour co-incubation compared to the growth of GBS alone. The presence of GBS did not significantly change the overall growth of CAF2-FR and *NRG1^OEX^-iRFP* in RPMI media (Supplemental Figure 5).

**Figure 4:**
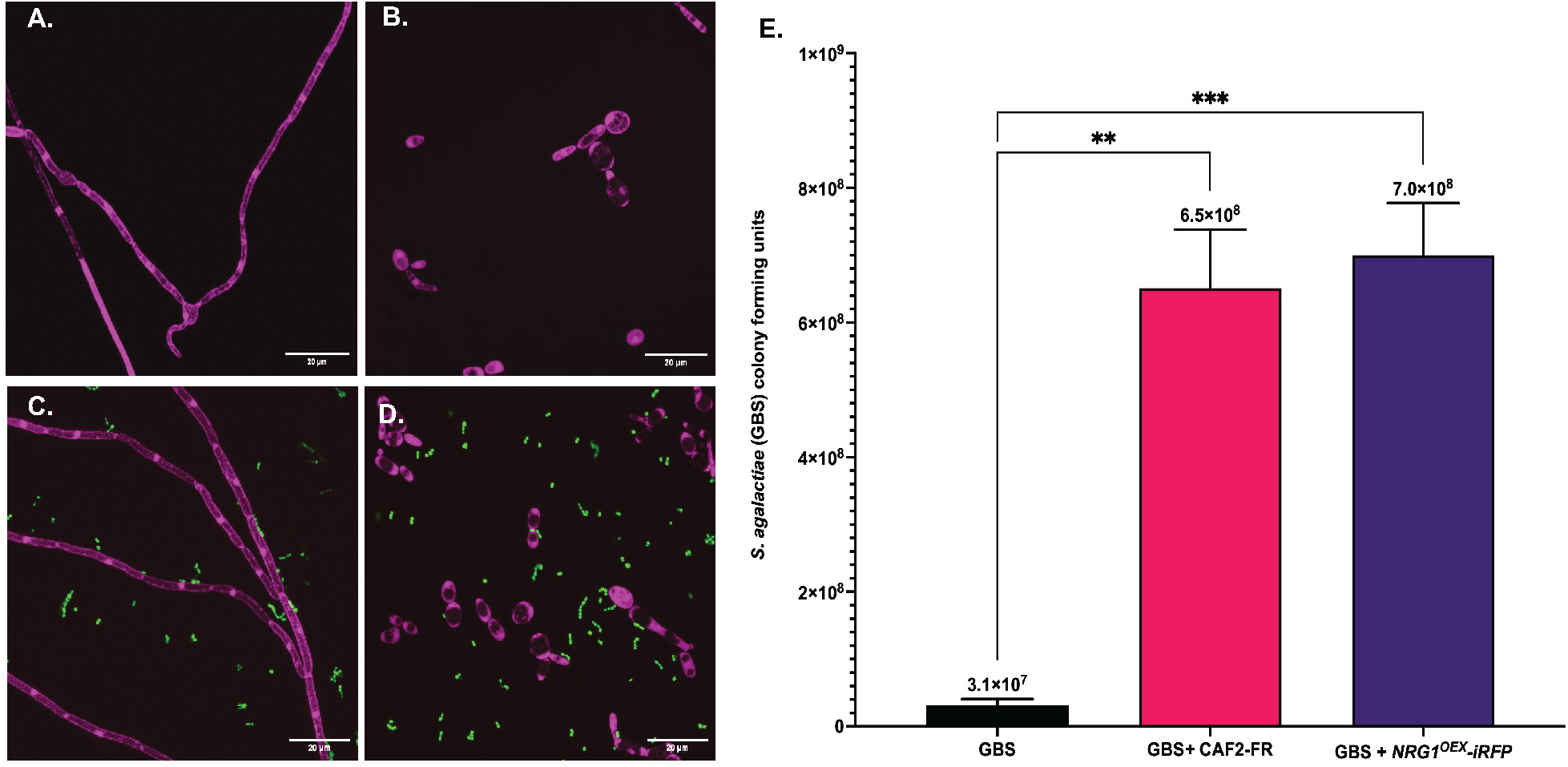
*C. albicans* hyphal formation is not required for GBS increased growth. GBS 515 was co-cultured with either a strain of *C. albicans NRG1^OEX^-iRFP* (yeast locked) or the parent strain that is freely able to hyphae, CAF2-FR for 24 hours in nutrient poor media. Fluorescent images were taken at 1000X magnification using a point-scanning confocal microscope (RFP = *C. albicans*; GFP = GBS) **A.** CAF2-FR following 24 hours of incubation in nutrient poor media. **B.** *NRG1^OEX^-iRFP* following 24 hours of incubation in nutrient poor media. **C.** Co-culture of GBS 515-GFP and CAF2-FR following 24 hour incubation in nutrient poor media. **D.** Co-culture of GBS 515-GFP and *NRG1^OEX^-iRFP* following 24 hours of incubation in nutrient poor media. **E.** Growth of GBS 515 following solo or co-culture with CAF2-FR or *NRG1^OEX^-iRFP*. Cultures were grown for 24 hours in nutrient poor media. Results are averaged from 4 biological replicates (n=4). Statistical significance was calculated using one-way ANOVA with Dunnett’s multiple comparison’s test, **p<0.005, ***p<0.0005.

### Physical interaction is not required for *C. albicans* to increase GBS growth *in vitro*

To determine if GBS and *C. albicans* need to have direct physical contact for the enhanced GBS growth observed during co-culture, we used a trans-well system in which the RPMI medium could be shared between the two organisms while keeping each organism separate. Both organisms were grown as detailed above for the co-culture experiments, followed by inoculation into either the well of a 6-well plate or into the 0.4 µM filter trans-well insert placed into the same well, which allows for media to be shared but no physical contact between the two organisms. Cultures were grown for 24 hours in a covered shaking water bath incubator with the covered plates attached to a rack held above the water level. This scenario served to replicate the growth conditions for the co-cultures described above. After 24 hours growth, GBS grown alone showed a very similar amount of growth as cultures grown for 24 hours in Figure 1 (Figure 5A). Similarly, when GBS and *C. albicans* were grown together in the same well, GBS showed a significant increase (p<0.0001) in growth as was shown for 24 hours in Figure 1 (Figure 5A). Surprisingly, we observed a significantly higher total growth (p>0.0001) when GBS and *C. albicans* shared the same media but were not in physical contact using the trans-well insert (Figure 5A). To determine if shared media in the absence of physical contact affects the growth of *C. albicans,* we measured growth at 24 hours under the same 3 conditions. Again, *C. albicans* solo growth was almost identical as in Figure 1 at 24 hours (Figure 5B). In contrast, we saw a decrease in growth of *C. albicans* when grown in the same well as GBS and a significant increase (p>0.0001) in growth in *C. albicans* when the organisms were separated by the trans-well insert but shared the same media (Figure 5B). This suggests that physical contact between GBS and *C. albicans* may inhibit growth of *C. albicans* by an unknown mechanism.

**Figure 5:**
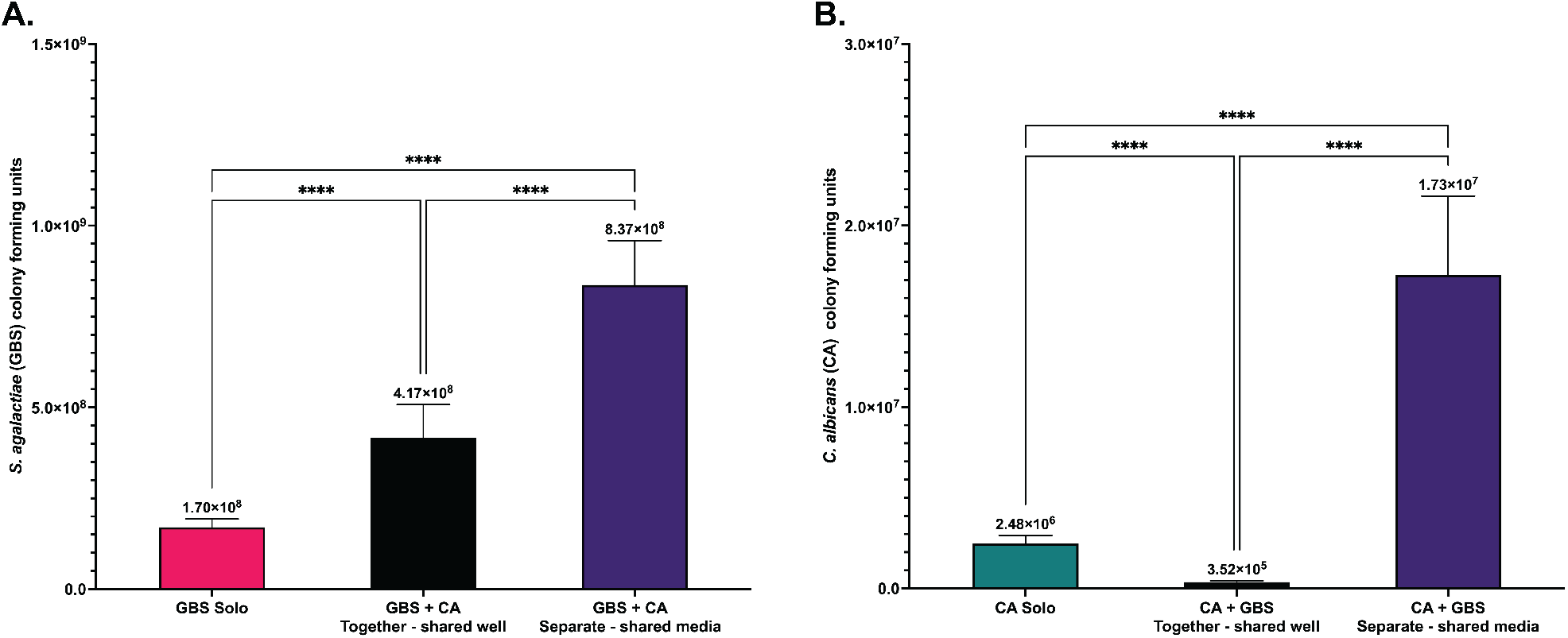
Physical interaction is not required for increased GBS growth in co-culture *in vitro*. **A.** Colony growth of GBS following solo or co-culture with *C. albicans* at 24 hours post culture in nutrient poor media grown together in a shared well or physically separated using a transwell with shared media. **B.** Colony growth of *C. albicans* following solo and co-cultured with GBS at 24 hours post culture in nutrient poor media grown together in a shared well poor physically separated using a transwell with shared media. Results above are from 3 combined experiments, with experiments performed in triplicate. Statistical significance was calculated by using Brown-Forsythe and Welch ANOVA, followed by Dunnett’s T3 multiple comparison test., ****p<0.0001.

### GBS antibiotic susceptibility is decreased when co-cultured with *C. albicans in vitro*

Antibiotic resistant GBS is a major clinical concern for pregnant women, who have the risk of passing the bacteria to their offspring in utero or during delivery, potentially leading to life threatening bacterial infections in their newborns. GBS strains are showing increased rates of resistance to both erythromycin and clindamycin clinically, which is a concern for people allergic to beta lactam antibiotics who rely on these antibiotics as alternative therapies (52–54).

Previous work demonstrated that *C. albicans* is able to enhance antibiotic tolerance of other Gram positive or negative bacteria including *Staphylococcus aureus,* and *S. gordonii* and *Pseudomonas aeruginosa* (19, 55–57). To investigate if the presence of *C. albicans* can affect antibiotic susceptibility of GBS during co-culture, both organisms were cultured together with either erythromycin or clindamycin. The growth of *C. albican*s was not significantly altered in solo or co-cultures with GBS treated with either antibiotic, indicating that these antibiotics do not affect the growth of *C. albicans* (Figures 6B & 6D). Erythromycin and clindamycin were still effective at decreasing the survival of GBS in both solo and co-cultures with *C. albican*s compared to cultures grown without the antibiotic (Figures 6A & 6C). However, co-cultures of GBS and *C. albicans* have significantly (p=0.002 and p=0.01, respectively) more GBS cells recovered following antibiotic treatment compared to solo cultures of GBS treated with erythromycin and clindamycin (Figures 6A & 6C). This indicates that *C. albicans* can alter erythromycin and clindamycin effectiveness against GBS *in vitro*.

**Figure 6:**
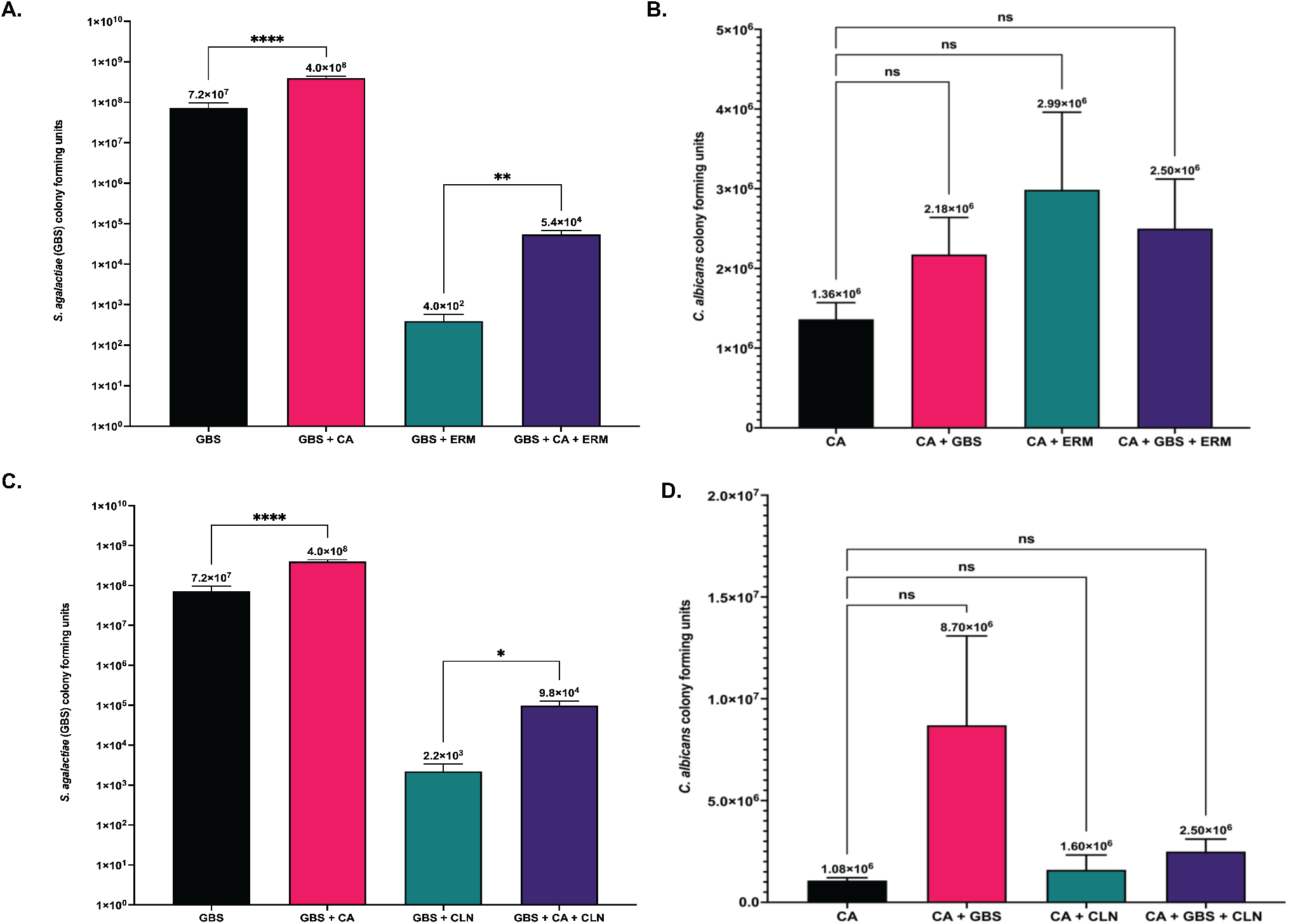
Antibiotic effectiveness against GBS is reduced in the presence of *C. albicans*. **A.** GBS growth following solo or co-culture with *C. albicans* treated with 2µg/mL erythromycin. **B.** *C. albicans* growth following solo or co-culture with GBS treated with 2µg/mL erythromycin. **C.** GBS growth following solo or co-culture with *C. albicans* treated with 2µg/mL clindamycin. **D.** *C. albicans* growth following solo or co-culture with GBS treated with 2µg/mL clindamycin. Experiments investigating treatment of cultures with erythromycin were performed 6 times (n=6). Experiments investigating treatment of cultures with clindamycin were performed 5 times (n=5). Results above are from combined experiments, and statistical significance was calculated by unpaired two-tailed student’s t test for A & C, and One-Way ANOVA with Dunnett’s multiple comparison’s test for B & D, *p<0.05, **p<0.005.

To determine if hyphal formation was required for the observed decreased susceptibility of GBS to antibiotics during co-culture with *C. albicans*, we used the *C. albicans* yeast-locked strain, *NRG1^OEX^-iRFP,* in co-cultures with GBS treated with and without antibiotics. Results revealed that hyphal formation does appear to help promote this effect, but is not completely necessary, as GBS still shows some increased growth in the presence of antibiotics when co-cultured with the yeast-locked strain (Supplemental Figure 6). Antibiotic treatment had no effect on the growth of *NRG1^OEX^-iRFP* (data not shown).

The ability of *C. albicans* to synergize with GBS to enhance its antifungal tolerance was also investigated, as the effectiveness of certain antifungals against *C. albicans* has previously been shown to be altered in the presence of other bacterial strains including *P. aeruginosa* (16). To investigate if antifungal effectiveness can be altered for *C. albicans* when cultured with GBS, growth was determined for both organisms when grown together and separately in RPMI media for 24 hours with either fluconazole or nystatin antifungals. Results indicated that the effectiveness of fluconazole and nystatin against *C. albicans* was not significantly different in solo vs co-cultures with GBS (Figure 7A & 7C), indicating that GBS does not affect the ability of *C. albicans* to tolerate these antifungals. While GBS growth was not significantly affected by any of the antifungals tested in solo or co-cultures with *C. albicans*, the amount of GBS recovered from co-cultures when *C. albicans* was treated with fluconazole or nystatin was slightly reduced compared to untreated co-cultures, indicating that the ability of GBS growth to be enhanced by *C. albicans* can be altered when *C. albicans* is treated with certain antifungals (Figure 7B & 7D).

**Figure 7:**
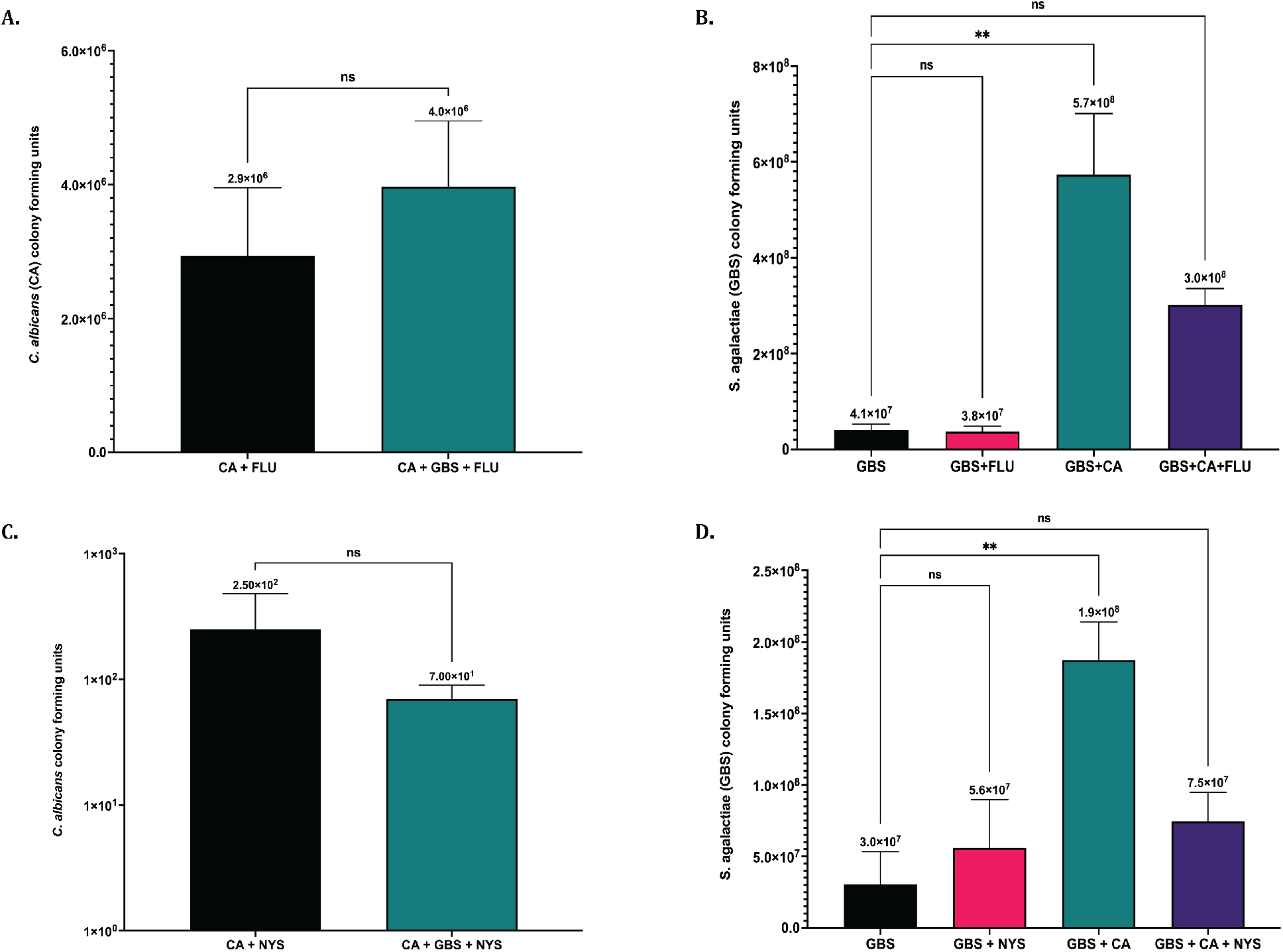
Antifungal effectiveness against *C. albicans* is not altered in the presence of GBS. **A.** *C. albicans* growth following solo or co-culture with GBS treated with 25µg/mL fluconazole. **B.** GBS growth following solo or co-culture with GBS treated with 25µg/mL fluconazole. **C.** *C. albicans* growth following solo or co-culture with GBS treated with 5µg/mL nystatin. **D.** GBS growth following solo or co-culture with GBS treated with 5µg/mL nystatin. Fluconazole experiments were performed in triplicate (n=3), and Nystatin experiments were performed 4 times (n=4). Statistical significance was calculated by unpaired two-tailed student’s t test for A & C, and One-Way ANOVA with Dunnett’s multiple comparison’s test for B & D. *p<0.05, **p<0.005, ****p<0.00005.

### Route of infection plays a role in virulence in *in vivo* co-infections with GBS and *C. albicans*

Data presented above revealed that interactions between GBS and *C. albicans* result in enhanced growth *in vitro.* Therefore, it was important to investigate if interactions between the two organisms can alter their virulence *in vivo*, as previous research had indicated that *C. albicans* virulence can be directly influenced by the presence of other microbes (58). The virulence of a co-infection compared to a solo infection of either GBS or *C. albicans* was investigated using a zebrafish infectious disease model. Zebrafish are well established as infection models for both streptococcal *spp.* and *Candida spp.* infections (59, 60). Injection into the otic vesicle (OV) was used as a “localized” infection model as the organisms are inoculated into an enclosed cavity that is devoid of leukocytes prior to infection. For solo infections, 40 colony forming units (cfu) of GBS or 40 cfu of *C. albicans* was microinjected into the OV of 2 days post fertilization (dpf) zebrafish larvae. For co-infections, half the dose of both organisms (20 cfu of GBS and 20 cfu of *C. albicans* for a total of 40 cfu) was injected into the OV. Injections of sterile media (5% PVP in PBS) were used as a negative control. Following injection, fish were monitored for survival every 24 hours for 72 hours total. When *C. albicans* was injected alone into the OV almost no death was observed (Figure 8). Zebrafish larvae infected with GBS alone at 40 cfu had a survival rate of ∼55% at 48 hours post infection (hpi). However, zebrafish co-infected with both GBS and *C. albicans* at half the dose of single infections showed significantly (p=0.002 and p=<0.0001, respectively) higher mortality rates by 48 hpi compared to solo infections with either GBS or *C. albicans* alone (Figure 8), with roughly 30% of the zebrafish surviving past 48 hours (Figure 8).

**Figure 8:**
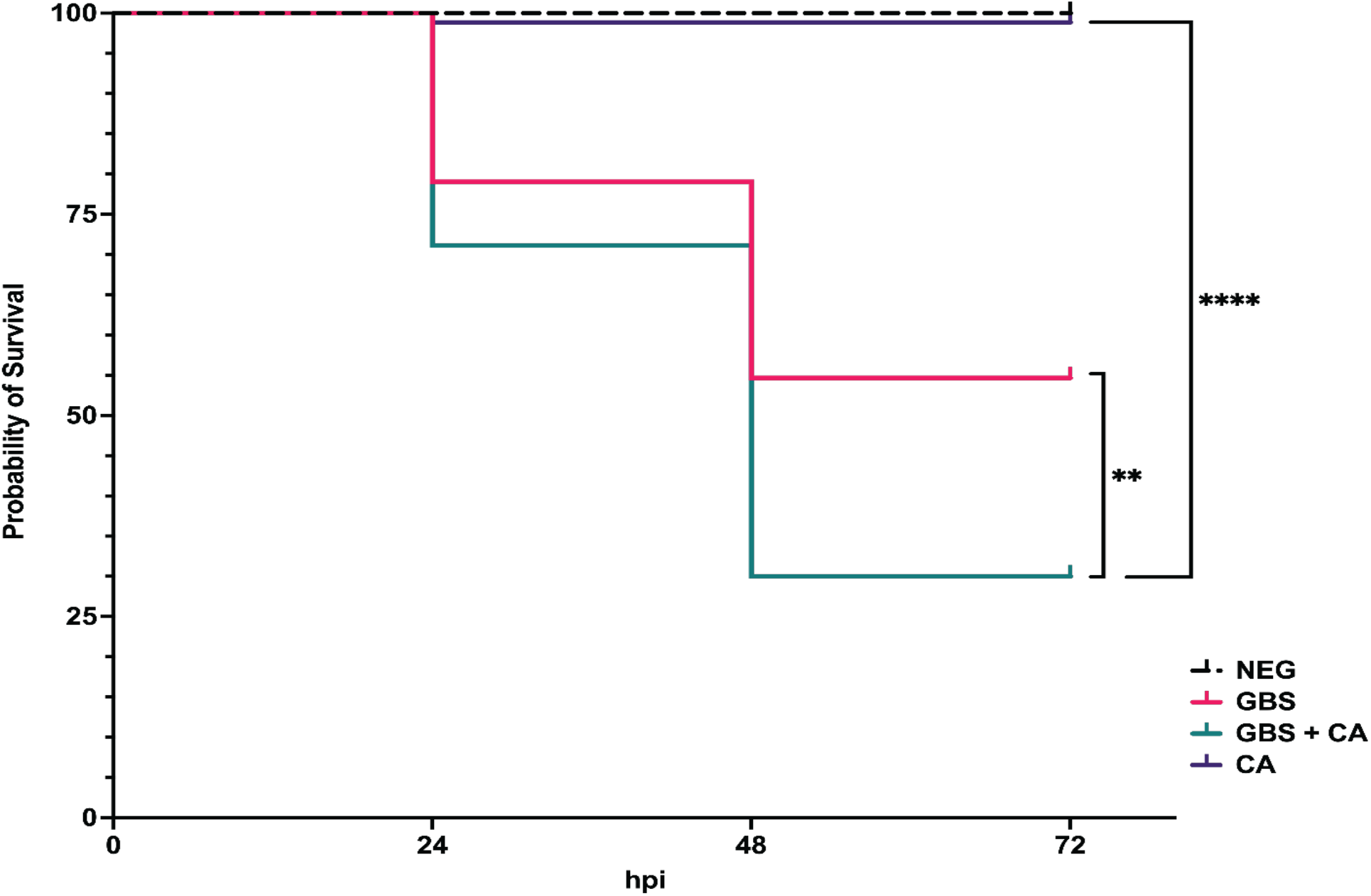
*C. albicans* and GBS synergize *in vivo* to increase virulence. Otic vesicle injection of solo or co-infections of GBS and *C. albicans* into 2dpf larval zebrafish. Solo injections had either 40 cfu of GBS 515 or 40 cfu of *C. albicans.* Co-infections had 20 cfu of GBS and 20 cfu of *C. albicans* (totaling 40 cfu) together. Negative control was 1 nL of 5% PVP in PBS. Infections were replicated 4 times with 20 fish per experimental condition (n=80). Above results are from combined experiments. Statistical significance of Kaplan-Meier survival curves was calculated using a log rank (Mantel-Cox) test. **p<0.05, ****p<0.00005.

To investigate if the bacterial and fungal burden in larval zebrafish is increased in co-infections of GBS and *C. albicans*, 2dpf zebrafish were injected into the OV with either a streptomycin resistant GBS strain or *C. albicans* individually, or a co-infection of both pathogens. In these assays, the same inoculum concentration was used for both solo and co-infections in order to accurately compare organism burden between the two conditions (see Materials and Methods). After 24 hpi zebrafish were euthanized, homogenized, and bacterial and fungal burden was determined by plating serial dilutions of the homogenized zebrafish onto selective plates. When zebrafish were injected with the same initial dose of GBS and *C. albicans* in co-infections compared to the dose injected with solo infections, co-infections led to a significantly (p=0.03) higher GBS bacterial burden in larval zebrafish at 24 hpi compared to solo infections of GBS (Figure 9A), indicating that *C. albicans* can enhance the amount of GBS recovered from infected fish in localized infections (Figure 9A). Fungal burden of *C. albicans* at 24 hours was not significantly different between solo and co-infections (Figure 9B). The average fungal burden of the larval zebrafish at 24hpi was less than the original inoculum, suggesting the immune system was able to clear *C. albicans* from the OV at this dose.

**Figure 9:**
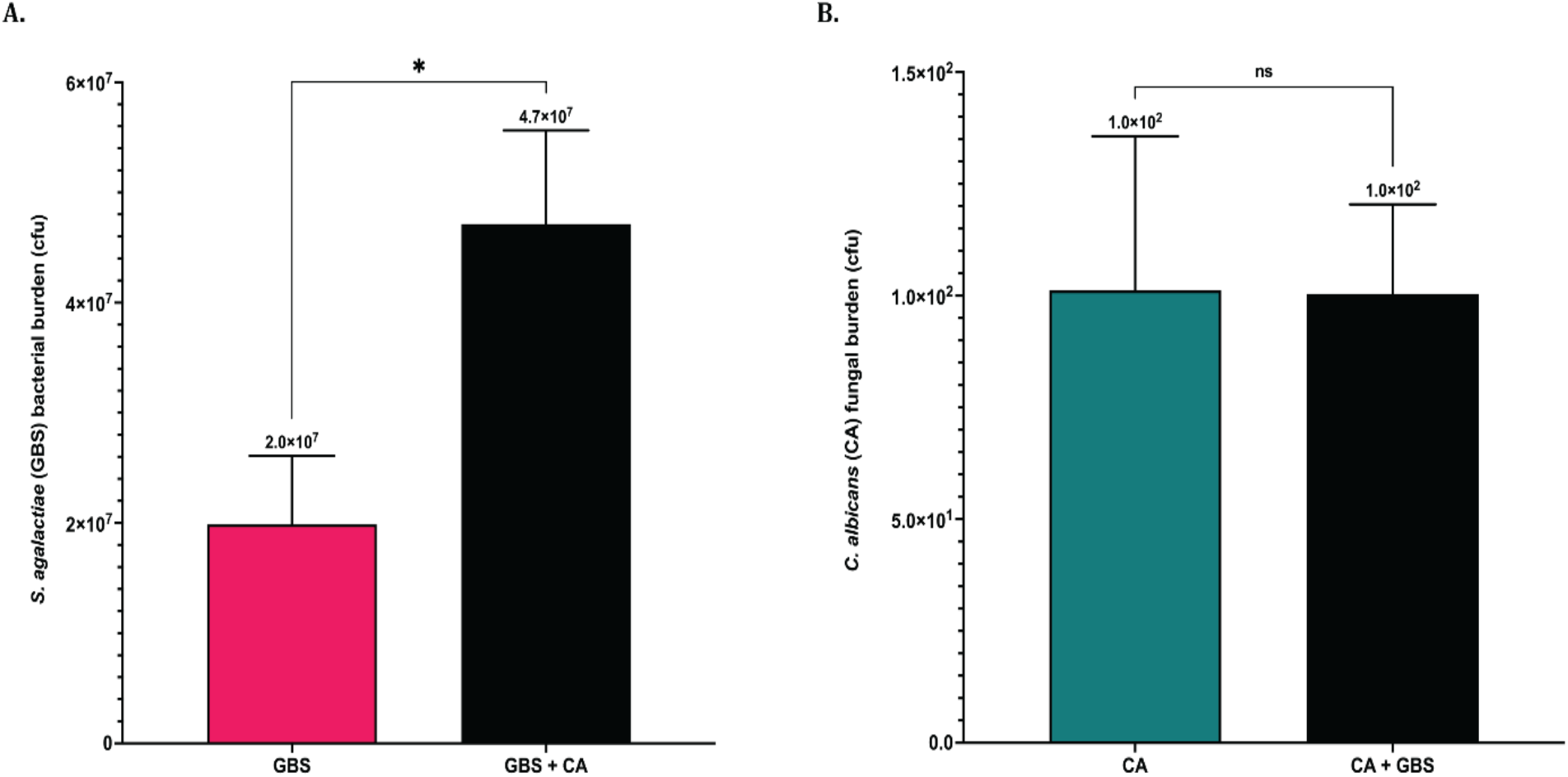
Localized co-infection with *C. albicans* increases GBS burden *in vivo* when injecting with the same initial GBS dose. The bacterial and fungal burden in infected zebrafish was calculated following otic vesicle injection of solo or co-infections of GBS and *C. albicans* into 2dpf larval zebrafish. Solo injections had either 40 cfu of GBS 515 or 40 cfu of *C. albicans.* Co-infections had 40 cfu of GBS and 40 cfu of *C. albicans* (totaling 80 cfu) together **A.** The average GBS bacterial burden in zebrafish 24 hours following localized solo or co-infection with *C. albicans* when fish were injected with the same starting dose of GBS (40 cfu) **B.** The average *C. albicans* fungal burden of zebrafish 24 hours following localized solo or co-infection with GBS when fish were injected with the same starting dose of GBS (40cfu). Experiments were performed 5 times (n=5). Above results are from combined experiments, with statistical significance calculated using an unpaired two-tailed student’s t test *p<0.05.

While infections centralized in the vaginal tract are considered localized, both of these pathogens have the capability of disseminating into tissues, potentially leading to severe bloodstream infections with high mortality rates (61, 62). Yolk sac injection of the zebrafish larvae with either of these pathogens most often leads to a systemic infection. Therefore, to investigate if co-infections of these pathogens are more virulent than solo infections of either pathogen, an infection route that most often leads to a systemic infection was used. Systemic infections in zebrafish were performed by injecting either GBS or *C. albicans* alone, or half of the dose of both pathogens (resulting in same overall dose as solo injections) into the yolk sac of 2 dpf larval zebrafish and monitoring survival for 72 hpi. This results in a bloodstream infection with no localized infection in the yolk sac. At 72 hpi results showed that there were no significant differences in mortality rates between solo infections of GBS and *C. albicans* and co-infections with both pathogens (Figure 10A). However, it should be noted that co-infections where half the bacterial and fungal load of GBS and *C. albican*s than solo infections, but were just as virulent as the solo infections, indicating that interactions between these two pathogens may promote virulence for these pathogens (Figure 10A).

**Figure 10:**
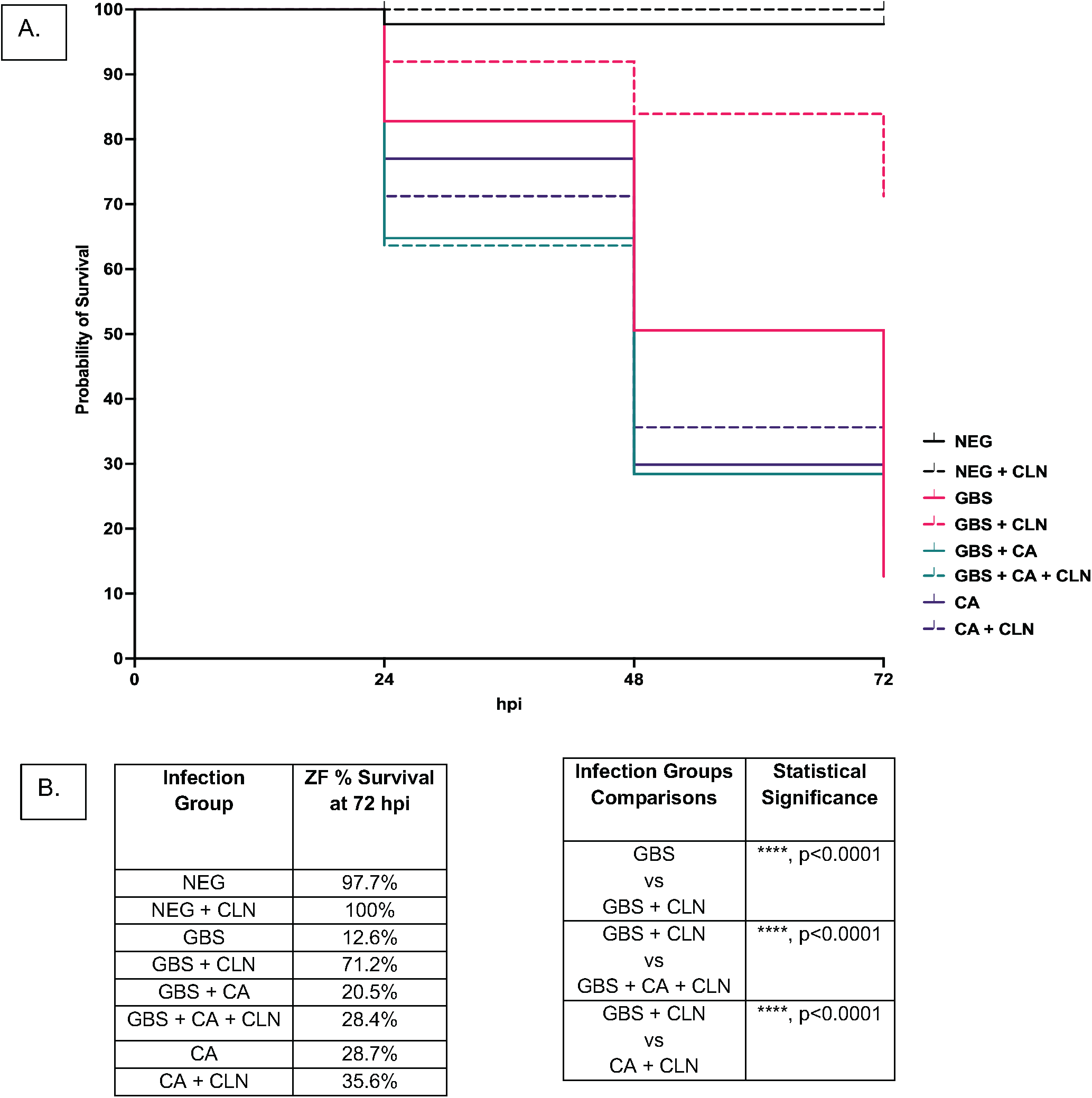
The presence of *C. albicans* decreases effectiveness of clindamycin against GBS *in vivo.* Solo or co-infections of GBS and *C. albicans* using a yolk sac injection method on 2dpf zebrafish larvae. For solo infections zebrafish were injected with either 20 cfu of GBS 515 or 20 cfu of *C. albicans.* For co-infections zebrafish were injected with 10 cfu of GBS and 10 cfu of *C. albicans* (totaling 20 cfu) together. The negative control was 1 nL of 5% PVP in PBS. Experiments were replicated 3 times with 20 fish per experimental condition. Graph shows pooled date of 3 experimental replicates **A.** Zebrafish survival of systemic solo and co-infection of GBS and *C. albicans* untreated or treated with 6µg/mL dose of clindamycin in the tank water of the fish at the time of initial infection. **B.** Tables describing zebrafish survival and infection group comparisons where differences in survival percentage were found to be statistically significant for systemic solo and co-infections of GBS and *C. albicans* untreated or treated with 6µg/mL clindamycin Statistical significance of Kaplan-Meier survival curves were calculated using a log rank (Mantel-Cox) test. ****p<0.0001.

### GBS is less susceptible to antibiotic treatment during systemic co-infections *in vivo*

Antibiotic treatment effectiveness for *in vivo* co-infections between GBS and *C. albicans* have not been studied previously. *In vitro* results reported above (Figure 6) indicated that when GBS is co-cultured with *C. albicans* in RPMI media that GBS is less susceptible to the antibiotics erythromycin and clindamycin. To determine if systemic co-infections of GBS and *C. albicans* are less susceptible to the antibiotic clindamycin, a survival assay was performed, and the same dose of clindamycin was administered to the water of the infected zebrafish for all infection groups directly following inoculation. Survival was monitored every 24 hours for 72 hours total. Results showed that the zebrafish mortality rate of solo infections of GBS treated with clindamycin were significantly (p=<0.0001) decreased by ∼59% compared to untreated solo GBS infections. However, co-infections of GBS and *C. albicans* treated with clindamycin were just as virulent as untreated co-infections, with less than an 8% reduction in mortality following treatment, indicating that GBS are less susceptible to the antibiotic clindamycin *in vivo* during a co-infection with *C. albicans*.

To determine if the hyphal form of *C. albicans* is required for the loss of susceptibility of GBS to clindamycin *in vivo*, infections were repeated as above with the yeast-locked *C. albicans* strain *NRG1^OEX^-iRFP.* The results confirmed that the clindamycin-treated systemic co-infections with GBS and yeast-locked *C. albicans* strain *NRG1^OEX^-iRFP* were significantly more virulent than clindamycin treated GBS solo infections (p=0.02), indicating that *C. albicans* does not need to be able to form hyphal filaments to reduce the susceptibility of GBS during infection *in vivo* (Supplemental Figure 7). Interestingly, untreated and clindamycin treated co-infections of GBS and *C. albicans* did not significantly enhance the GBS recovered from co-infected fish compared to solo GBS infected fish (Supplemental figure 8). The combined results above indicate that the ability of *C. albicans* to decrease the susceptibility of GBS to the antibiotic clindamycin is not due to increased bacterial burden during infection.

## Discussion

The role of polymicrobial interactions on treatment and infection outcomes in the vaginal tract has been understudied despite increased knowledge about the influence of the microflora on disease in other tissue environments. The vaginal tract is capable of rapidly changing the composition of its microbiome, while being a challenging tissue environment to colonize due to its acidic mucosa (63–66). While the vaginal tract is highly populated with the beneficial bacteria *Lactobacillus* in healthy individuals, the overgrowth of other microbes that can colonize this environment following disruptions to the lactobacilli population can cause serious health issues in patients (63–66). Two of these microbes, *C. albicans* and GBS, are especially dangerous in immunocompromised patients, such as newborns, patients with chronic diseases, the elderly, and pregnant women (2–5). GBS and *C. albicans* can synergize in colonization of the bladder and vaginal tract (39–41, 43), but not much was known prior to this study about the effects of interactions between these two pathogens beyond their ability to co-associate in these specific tissue environments. In this study, we found that co-association between GBS and *C. albicans* drives changes in growth and survival of GBS, especially in the context of antibiotic treatment.

In nutrient-limiting but not rich media, we find that *C. albicans* significantly improves GBS survival and growth. Both macro-and micro-nutrients such as iron, oxygen, and carbon-source drive important interactions between *C. albicans* and other bacterial species, including antifungal susceptibility, cell wall epitope exposure, and morphology (16, 34, 67–72). The same is true for GBS. Nutrient availability impacts gene expression for both *C. albicans* and GBS, and the gene expression of both would likely influence the way these microbes interact with one another *in vitro* and *in vivo*(73–77). These nutrient-dependent interactions are important to consider in concert with the fact that nutrient availability in vaginal tract secretions is regulated by pregnancy, poor diet, hormones and stress (63–66). In addition, we found that physical contact between the two organisms was not required for the enhanced growth of GBS when the two organisms shared the same media. This suggests that there is some molecule or nutrient that is released into the media that can be utilized by GBS for growth. Our trans-well data also suggests that physical contact between GBS and *C. albicans* may inhibit growth of *C. albicans* when grown in a nutrient limited environment. We also observed a surprising increase in growth of *C. albicans* when it shared media with GBS, but physical contact was prohibited, showing a ∼7-fold increase in growth over *C. albicans* is grown alone. Whether this is nutrient based or a molecule that acts as a signal for increased growth is unknown at this point. The ability of GBS to co-associate with *C. albicans* may be nutrient and environmentally dependent, which may explain why certain patients are at a higher risk of infection and serious complications by these pathogens.

*C. albicans* cell morphology has been shown to affect interactions with GBS. *C. albicans* is polymorphic, can grow in three distinct cellular morphologies (yeast, pseudohyphae, and hyphae) and can alter its morphology based on environmental factors such as nutrient availability, temperature, stress, and fluctuations in pH (78–81). We found that *C. albicans* hyphal growth is not inhibited by the presence of GBS and does not affect its ability to enhance GBS growth. This suggests that the ability of GBS to inhibit hyphal growth may be environmentally influenced or strain specific. Previous studies of interactions between GBS and *C. albicans* have had conflicting conclusions regarding the role of *C. albicans* cellular morphology in the co-association of these two microbes (41–43). Some studies have indicated that the co-association of GBS and *C. albicans* depends on the expression of the hyphal-specific invasin Als3p (41, 43, 82). Als3p is also responsible for the synergy of *C. albicans* and GBS in their colonization of the bladder of mice (43, 82). On the other hand, GBS can inhibit *C. albicans* hyphal growth by decreasing the expression of *EFG1*-induced Hwp1, a hyphal adhesion protein required for biofilm production (42). Taken together, our results indicate that hyphal morphology and Als3p are only responsible for some aspects of *C. albicans*-GBS interaction, which may rely on multiple factors such as environmental stressors, specific strain traits, and nutrient availability.

This study aimed to investigate how interactions between *C. albicans* and GBS can influence treatment effectiveness, as antimicrobial resistance by both pathogens can have serious health implications clinically Polymicrobial interactions between *C. albicans* and GBS significantly altered the effectiveness of erythromycin and clindamycin *in vitro* against GBS compared to GBS cultured alone with these treatments. However, no significant changes in treatment effectiveness against *C. albicans* by multiple antifungal treatments were observed during co-culture with GBS. While vaccines against GBS have been in development for several years, there is no vaccine commercially available against GBS, and no alternative treatment options for these infections are available outside of antibiotic use. The main antibiotic used for both intrapartum antibiotic prophylaxis (IAP) administration and to treat GBS invasive infection is penicillin, a beta-lactam antibiotic considered to be highly effective against GBS (6). Erythromycin and clindamycin are two antibiotics prescribed to patients who cannot be treated with penicillin due to allergies to beta-lactam antibiotics. Clinically, GBS strains are showing rising resistance to both erythromycin and clindamycin, with clindamycin-resistant Group B *Streptococcus* strains specifically being implicated as a concerning threat by the CDC (83). Erythromycin and clindamycin belong to different families of antibiotics (macrolide and lincomycin, respectively) but have similar mechanisms of action that target the 50s ribosomal subunit, preventing protein synthesis (84). Commonly, GBS clinical strains show cross-resistance to erythromycin and clindamycin, leading to both being unavailable treatment options for many patients (84). Previous research has shown that vaginal candidiasis caused by *C. albicans* can occur following the use of antibiotics (22). *C. albicans* can also become less susceptible to amphotericin B when exposed to erythromycin, indicating that antibiotics can influence the behavior of *C. albican*s (85). The risk of *C. albicans* co-colonization with other multidrug resistant organisms has also been shown to be increased in hospital patients previously treated with macrolide antibiotics, highlighting the influence of antibiotics on *C. albicans* (86). Our results show that the antibiotics erythromycin and clindamycin were significantly less effective against GBS *in vitro* when co-cultured with *C. albicans* in nutrient poor media in comparison to solo GBS cultures. These results indicate that these antibiotics may not be the best option for patients colonized with both *C. albicans* and GBS, as they are not only less effective against GBS, but can potentially alter the susceptibility of *C. albicans* to certain antifungals following exposure to these antibiotics.

We found that co-infections of GBS and *C. albicans* are significantly more virulent than solo infections of either pathogen in the larval zebrafish otic vesicle localized infection model. This is consistent with enhanced *C. albicans*-*Streptococcus* pathogenesis in rodent co-infection models, with increased tissue damage, pathogen burden and tooth carious lesions (33–38, 42, 43)(42, 43). However, how GBS and *C. albican*s co-infections alter infection outcomes in comparison to solo infections in either a localized or a systemic infection is not well understood. Our study utilized larval zebrafish as an *in vivo* infection model for studying co-infection with GBS and *C. albicans*, because there are well-characterized zebrafish models for both pathogens and this simpler transparent vertebrate offers several advantages over mice (59, 60, 87, 88). For instance, zebrafish are transparent, offer multiple routes of infection, and enable treatment administration in the tank water. By utilizing both *in vitro* and *in vivo* experimental models, we can better understand how interactions between these organisms can alter virulence and treatment effectiveness, and these findings can be used to further investigate how treatment effectiveness is altered by GBS and *C. albicans* interactions in a mammalian host (59, 60).

In contrast to the localized infection model, there were no significant enhancements in virulence for GBS and *C. albicans* yolk sac co-infections. This may be because disseminated infections are severe for both individual infections. It is also possible that there is less contact between the organisms in disseminated, as compared to the localized otic vesicle infection, resulting in less synergy. Not surprisingly, clindamycin treatment rescued GBS solo infections. However, clindamycin failed to rescue co-infections with *C. albicans* and GBS, indicating that the presence of *C. albicans* makes the antibiotic clindamycin less effective against GBS following co-infection in larval zebrafish *in vivo.* This matches our *in vitro* data that clindamycin is less effective against GBS when *C. albicans* is present. While the treatment effectiveness of antibiotics against GBS in the presence of *C. albicans* has not been researched prior to this study, it has been shown previously that mixed biofilms of *C. albicans* and other bacterial species are less susceptible to antimicrobial treatments (19, 57, 89). Our results suggest that clindamycin may not be a suitable alternative antibiotic against GBS for patients that are also colonized with *C. albicans*, highlighting the importance of knowing the microbial composition of the vaginal tract for patients, especially expecting mothers.

The results from this study demonstrate that co-colonization between GBS and *C. albicans* can alter microbial growth, treatment effectiveness, and virulence of infections for both pathogens. Overall, these results help further define the ability of GBS and *C. albicans* to co-associate in specific conditions and highlight important clinical issues for treatment effectiveness and infection outcomes for both pathogens.

## Materials and Methods

### Bacterial strains and growth conditions

The *Streptococcus agalactiae* strain GBS 515 was used in all experiments unless otherwise noted (Bacterial strains listed in Table 1). GBS 515 is a serotype 1a, ST-23 human clinical isolate from the blood of a patient diagnosed with neonatal septicemia and was generously provided by M.R. Wessels (90). For all experiments using GBS, the bacterium was inoculated in Todd-Hewitt medium (Acumedia) supplemented with 0.2% yeast extract (THY) in sealed conical tubes and grown statically at 37°C overnight. For selection of GBS after co-culture experiments, cultures are serially diluted and plated on Strep B ChromoSelect Selective Agar Base (Millipore Sigma) agar plates.

**Table 1:**
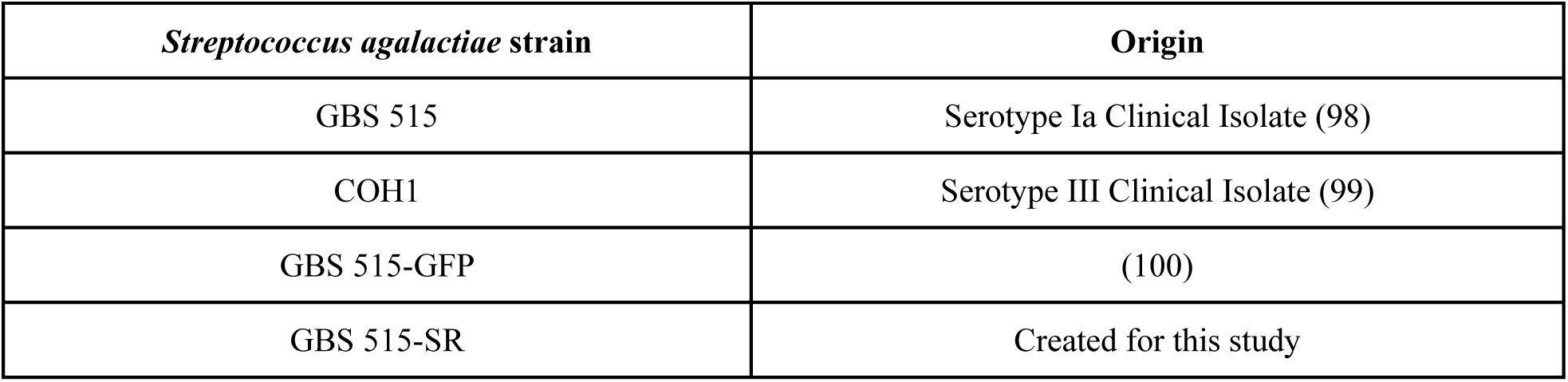
*Streptococcus agalactiae* strains in this study

### Fungal strains and growth conditions

The fungal strain of *Candida albicans* (*C. albicans*) used for co-culturing experiments with and without specific antifungals or specific antibiotics was SC5314-NEON unless otherwise noted (Fungal strains listed in Table 2). *C. albicans* was inoculated onto a THY agar plate for individual colonies and grown overnight at 37°C aerobically. For liquid overnight cultures, a single colony of *C. albicans* was selected from an agar plate, inoculated into THY media, and grown overnight with shaking aerobically in glass culture tubes at either 30°C or 37°C as noted in the experiment. For selection of *C. albicans* after co-culture experiments, cultures are serially diluted and plated onto THY plates supplemented with 2µg/mL ampicillin (Millipore Sigma).

**Table 2:**
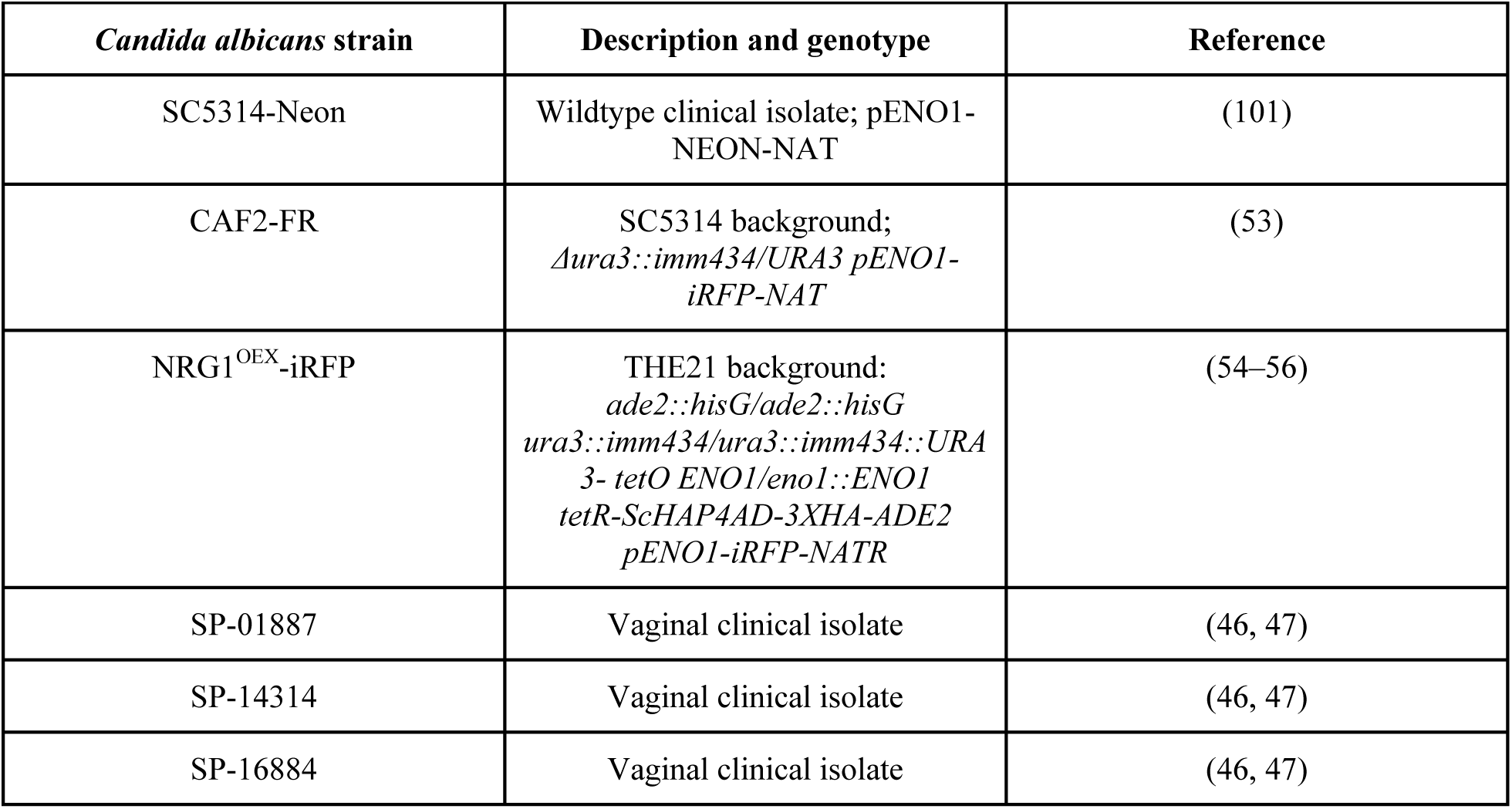
*Candida albicans* strains in this study

### Growth curve experiments

For growth curve experiments GBS 515 and SC5314-NEON were both grown overnight in THY media aerobically at 37°C. GBS 515 was grown statically overnight, while liquid cultures of SC5314-NEON were grown with shaking. Following 14 hr. overnight incubation, GBS 515 was sub-cultured into fresh THY media at a 1:100 dilution and subcultures were grown and normalized to an optical density (OD) 600 nm of 0.225 (mid log phase) prior to co-culturing. Prior to co-culturing, 1mL of the overnight culture of SC5314-NEON was added to a sterile Eppendorf tube, briefly vortexed, then centrifuged at 7000 RPM for 1 minute. Following centrifugation, the supernatant was removed, and the SC5314-NEON pellet was resuspended in 1 mL of sterile 1X PBS. This step was repeated two times, then following the last resuspension the concentration of SC5314-NEON was determined by measuring the absorbance of a 1:100 dilution of the SC5314-NEON in 1 mL of PBS at OD600 nm. Once concentrations of GBS 515 and SC5314-NEON were calculated, GBS 515 and SC5314-NEON were inoculated either together or in solo in 6 mL of nutrient rich (THY) or nutrient poor (RPMI 1640 with L-glutamine and 25mM HEPES) media at a concentration of 2 X 10^5^ cfu/ml per organism. Following culturing, serial dilutions of solo and co-cultures of GBS 515 and SC5314-NEON grown with shaking were plated once an hour for 6 hours totals on selective media plates and grown for 24 hours at 37°C. Following growth on selective media, individual colonies were enumerated to determine overall growth concentration of solo and co-cultures of GBS 515 and SC5314-NEON for each time point.

### Co-culture experiments with antimicrobials

For co-culture experiments with and without specific antimicrobials, GBS 515 and SC5314-NEON were both grown overnight in THY media aerobically at 37°C as described above for growth curve experiments. Once concentrations of GBS 515 and SC5314-NEON were calculated, GBS 515 and SC5314-NEON were inoculated either together or in solo in 4 mL of nutrient rich (THY) or nutrient poor (RPMI 1640 with L-glutamine and 25mM HEPES [LONZA]) media at a concentration of 2 X 10^5^ cfu/mL per organism at 37°C with shaking. Following culturing, serial dilutions of solo and co-cultures of GBS 515 and SC5314-NEON with or without supplemented antimicrobials were plated at either 6 hours or 24 hours post co-culture on selective media plates and grown for 24 hours aerobically at 37°C. When treated with fluconazole (Millipore Sigma) cultures were supplemented with a dose of 25 µg/mL. When treated with erythromycin (Millipore Sigma) tubes were supplemented with a dose of 2 µg/mL. When treated with clindamycin (Indofine Chemical Company, Inc, Hillsborough, NJ) tubes were supplemented with a dose of 2 µg/mL. To enumerate GBS in experimental cultures serial dilutions from solo and co-cultures that were untreated were plated on Strep B ChromoSelect Selective Agar Base (Millipore Sigma), grown overnight at 37°C and individual colonies were enumerated. GBS cultures treated with antibiotics erythromycin or clindamycin were plated on THY agar plates supplemented with 50 µg/mL Nystatin (Millipore Sigma), grown overnight at 37°C aerobically, and individual colonies were enumerated. To enumerate SC5314-NEON concentrations in experimental cultures serial dilutions were plated on THY agar plates supplemented with 2 µg/mL ampicillin (Millipore Sigma), grown overnight at 37°C and individual colonies were counted.

To confirm that co-culture results with GBS were not strain specific, GBS clinical isolate, serotype III CC-17 strain COH1 was co-cultured without antimicrobial treatment as described above. Following 24 hours of culturing, the growth of solo cultures of COH1 and SC5314-NEON, as well as co-cultures of both organisms were calculated following the enumeration of individual colonies on selective plates following serial dilutions. COH1 and SC5314-NEON selective plates used for this experiment are the same as the selective plates described above.

### Co-culture experiments using transwell system

For co-culture experiments using a six-well plate and transwell system (ThinCerts, USA Scientific) GBS 515 and SC5314-NEON were grown overnight in THY media aerobically at 37°C as described above for growth curve experiments. Once the concentrations of GBS 515 and SC5314-NEON were calculated, each organism was inoculated individually into 4 mL of nutrient-poor media (RPMI 1640 with L-glutamine and 25 mM HEPES) at a concentration of 2 × 10^5^ cfu/mL per organism in individual wells of a six-well plate. For co-culture experiments, GBS 515 and SC5314-NEON were either inoculated together in a shared well or inoculated in a shared media setup, where SC5314-NEON was added to the well and GBS 515 was placed in a 0.4 μm pore transwell insert. The covered six-well plate was attached to a 50 mL test tube rack situated above the water line and incubated with shaking at 37°C for 24 hours.

Following the 24-hour incubation, serial dilutions of solo and co-cultures of GBS 515 and SC5314-NEON were plated on selective media plates. To enumerate GBS, serial dilutions from solo and co-cultures were plated on THY agar plates supplemented with 50 µg/mL Nystatin (Millipore Sigma), grown for 24 hours at 37°C aerobically and individual colonies were counted. To enumerate SC5314-NEON, serial dilutions were plated on THY agar plates supplemented with 2 µg/mL ampicillin (Millipore Sigma), grown for 24 hours at 37°C aerobically and individual colonies were counted. To confirm that each organism did not cross-contaminate between the well and transwell insert, samples were plated on selective media as controls. An undiluted sample of the SC5314-NEON from the well was plated on THY agar plates supplemented with 50 µg/mL Nystatin to identify any GBS contamination. An undiluted sample of the GBS from the transwell insert was plated on THY agar plates supplemented with 2 µg/mL ampicillin to identify any SC5314-NEON contamination. Growth on either of these plates would indicate cross contamination between the well and transwell. Trial experiments were performed with GBS and *C. albicans* in either the well or the transwell to determine if location made a difference in growth. No difference was observed in results depending on location (data not shown).

### Heat-killed experiments

For heat-killed *C. albicans* experiments, GBS 515 and SC5314-NEON solo cultures and co-cultures were prepared using the same co-culture procedure described above, but prior to adding SC5314-NEON to the cultures the *C. albicans* cells was boiled at 100°C for 15 minutes prior to introduction into nutrient poor media solo and co-cultures. Heat-killed *C. albicans* was also plated on THY agar supplemented with 2 µg/mL ampicillin to confirm that cells were no longer viable. Solo and co-cultures of GBS 515 and heat killed SC5314-NEON were grown for 24 hours at 37°C aerobically with shaking, and GBS 515 growth was calculated by plating serial dilutions of cultures on the same selective media plates as stated above.

### Hyphal *C. albicans* strain vs. Yeast-locked strain assay

For experiments investigating the role of hyphal growth on GBS growth rates, GBS 515 was grown statically for 14 hours in THY media at 37°C and then sub cultured at a 1:100 dilution into 5 mL of fresh THY media. Subcultures were grown to an optical density (OD) 600 nm of 0.225 (mid log phase) prior to co-culturing with *C. albicans*. *C. albicans* strains CAF2-FR (reference strain capable of hyphal formation) and *NRG1^OEX^-iRFP* (yeast locked) were grown at 37°C overnight aerobically with shaking. Following incubation at 37°C for 16 hours, 1 mL of overnight stock was briefly vortexed, spun down at 7000 RPM for 1 minute and washed 2 times with 1X PBS before measuring the absorbance at OD 600 nm to determine concentration of each strain. After the concentration of cultures of GBS, CAF2-FR, and *NRG1^OEX^-iRFP* were determined, culture tubes with 4mL of RPMI 1640 with L-glutamine and HEPES (Lonza) were inoculated with a 2X10^5^ cfu/ml concentration of 515 GBS, a 2X10⁵ concentration of CAF2-FR, a 2X10⁵ concentration of *NRG1^OEX^-iRFP*, or 2X10⁵ concentration of GBS and a 2X10⁵ concentration of *C. albicans*. Cultures were grown for 24 hours and enumerated by counting individual colonies following serial dilutions plated on selective plates.

To image GBS 515, CAF2-FR, or *NRG1^OEX^-iRFP* in solo or co-cultures, cultures were inoculated as described above and grown at 37°C with shaking for 24 hours. Following this growth period, 7 µL of each culture were added to a glass slide and covered with a 22 x 22 mm cover slip. After the cultures were mounted on the glass slides confocal images were taken using an Olympus IX-81 inverted microscope containing a FV-1000 laser scanning confocal system (Olympus, Waltham, MA). The fluorescent proteins EGFP (488nm/505 to 525 nm excitation/emission) and Far-Red (635nm/ 655 to 755 nm excitation/emission) were detected using laser/optical filters with a 100X oil objective (NA,1.40). Images were taken and were processed using FluoView (Olympus, Waltham, MA). Image brightness was enhanced for visualization using ImageJ (90).

### Zebrafish care and maintenance

Adult zebrafish used for breeding were maintained at the University of Maine Zebrafish Facility at 29°C on recirculating systems. All zebrafish studies were approved by the University of Maine Institutional Animal Care and Use Committee (IACUC protocol #A2020-02-01). After collection, embryos were stored in petri dishes with sterilized water obtained from the recirculating system supplemented with 0.1% methylene blue. Embryos found to not be developing were removed after 24 hours post fertilization (hpf) to maintain cleanliness in petri dishes. Embryos were maintained at a temp of 29°C through development at a density of 50 embryos per dish. Following use in experiments, zebrafish were transferred to a 31°C incubator and were monitored for survival every 24 hours for 72 hours. Surviving zebrafish from experiments as well as zebrafish not used for experiments were humanely euthanized after 72 hours post experiment by an overdose of 0.64 µg/mL tricaine (Ethyl 3-aminobenzoate methanesulfonate (H_2_NC_6_H_4_CO_2_C_2_H_5_·CH_3_SO_3_H); Sigma-Aldrich). Experiments were conducted using the wild-type ZF1 strain.

### Zebrafish microinjections for localized infection

Larvae at ∼48 hpf were manually dechorionated and anesthetized in 0.32 mg/mL of tricaine prior to injection. Survival assays were performed by injecting either 1nL of GBS 515 at 4 X 10^7^ cfu/ml dose, 1nL of SC5314-NEON at 4 X 10^7^ cfu/ml dose, or 1nl of GBS at 2 X 10^7^ cfu/ml and SC5314-NEON at 2 X 10^7^ cfu/ml combined to equal an approximate dose to solo infections into the otic vesicle of larval zebrafish to simulate a localized infection (91). Mortality of the fish was determined by observation of a loss of heartbeat and no movement when gently probed on the tail. Survival of the zebrafish were monitored daily, and experiment was terminated at 72 hours. Surviving fish were euthanized with an overdose of tricaine (0.64 mg/mL) at the end of the experiment. Injection doses were confirmed by serial dilution and plating of the inoculum on THY agar plates and enumerating bacteria or fungi colonies.

### Zebrafish microinjections for systemic infection

Larvae at ∼48 hpf were manually dechorionated and anesthetized in 0.32 mg/mL of tricaine prior to injection. Survival assays were performed by injecting either 1 nL of GBS 515 at 2 X 10^7^ cfu/ml, 1 nL of SC5314-NEON at 2 X 10^7^ cfu/ml, or 1 nl of GBS at 1 X 10^7^ cfu/ml and SC5314-NEON at 1 X 10^7^ cfu/ml combined to equal an approximate dose equal to solo infections into the yolk sac of the larval zebrafish to simulate an infection as described previously (92). Mortality of the fish was determined by observation of a loss of heartbeat and no movement when gently probed on the tail. Survival of the zebrafish was monitored daily, and experiment was terminated at 72 hours. Surviving fish were euthanized with an overdose of tricaine (0.64 mg/mL) at the end of the experiment. Injection doses were confirmed by serial dilution and plating of the inoculum on THY agar plates and enumerating bacteria or fungi colonies.

To investigate if clindamycin effectiveness against GBS in solo vs co-infection is altered during systemic infection, clindamycin was added to the tank water of the zebrafish immediately after injection at a dose 6 µg/mL. Prior to these experiments a cytotoxicity screen was performed to determine if adding clindamycin to the tank water of zebrafish could cause harm to the fish by monitoring development in different doses of clindamycin over a 3-day period. Cytotoxicity was considered to be signs of deformity during early development of embryos including deformed or missing heads, hearts, or tails. No cytotoxicity was found for zebrafish treated with clindamycin at doses as high as 10 µg/mL (data not shown).

### Creation of GBS 515 Streptomycin resistant strain

To select for GBS 515 after injection into a zebrafish, a streptomycin resistant strain of GBS 515 was created (515 SR) following a previously described protocol (93). To spontaneously induce antibiotic resistance to streptomycin, overnight cultures of GBS 515 were grown in THY media with different concentrations of streptomycin, and then plated on THY agar plates supplemented with streptomycin. Spontaneous antibiotic resistance by GBS 515 to streptomycin was achieved after exposure to 250 µg/mL of streptomycin in liquid cultures and 250 µg/mL of streptomycin supplemented on agar plates. GBS 515 strain with streptomycin resistance was confirmed by streaking a liquid culture of the bacterium on an agar plate supplemented with 250 µg/mL of streptomycin, then performing a 16S ribosomal RNA (rRNA) gene polymerase chain reaction (PCR) specific for GBS. Multiple virulence assays were performed in zebrafish to confirm that the GBS 515 SR strain had the same virulence as the wild-type strain (Supplemental figure 9). However, it should be noted that the genome of this strain was not sequenced to determine that no secondary mutations were present.

### Bacterial and Fungal Burden Assay

Larvae at ∼48 hours post injection (hpi) were manually dechorionated and anesthetized using 0.32 mg/mL of tricaine prior to injection. Bacterial and fungal burden assay was performed by injecting either 1 nL of GBS (515 SR) at 4 X 10^7^, 1 nL of *C. albicans* (SC5314-NEON) at 4 X 10⁷ or 1 nL of GBS at 4 X 10^7^ and *C. albicans* at 4 X 10⁷ into the otic vesicle. Doses were confirmed by plating serial dilutions of inoculum on THY agar. At 24 hpi, 8 fish per infection group were euthanized by overdose of tricaine and homogenized using 200 µL of PBS and a pellet pestle (Thermo Scientific). Fish who were dead prior to euthanasia were not selected for homogenization. Following homogenization, the sample was plated on selective plates using serial dilution and approximate burden was calculated by enumeration of single colonies on selective plates. To calculate GBS bacterial burden, serial dilutions of the homogenized fish samples were plated on CNA agar with 250 µg/mL Streptomycin. To calculate *C. albicans* fungal burden samples were plated on Candida BCG agar (Difco) supplemented with Neomycin (500 µg/mL) and Ampicillin (10 µg/mL).

## Statistical analysis

All experiments were analyzed for significance using a two-tailed student’s t-test or one-way ANOVA with Dunnett’s multiple comparison’s test in Prism GraphPad unless otherwise noted in figure legends.

